# Idea Paper: Effects of gonad type and body mass on the time required for sex change in fishes

**DOI:** 10.1101/2021.08.29.458130

**Authors:** Soma Tokunaga, Tatsuru Kadota, Yuuki Y. Watanabe, Tetsuo Kuwamura, Yuuki Kawabata

## Abstract

Sex change is a well-known phenomenon in teleost fishes, and it takes several days to a few months depending on the species and direction of sex change. However, the underlying factors influencing the time required for sex change (*T*_*S*_) remain unclear. Given that the time for producing a new gonad largely determines *T*_*S*_, the gonad type (i.e., whether fish retain the gonad of opposite sex or not [delimited or non-delimited]) and metabolic rate are the ultimate determinants of *T*_*S*._ This study sought to test two hypotheses: (1) the delimited gonad shortens *T*_*S*_; and (2) *T*_*S*_ scales with mass^0.1–0.2^, because the metabolic scaling exponent (*β*) in fishes is 0.8–0.9 and biological times scale with mass^1−*β*^ in general. We compiled data on *T*_*S*_ for 12 female-to-male and 14 male-to-female sex-changing species from the literature. Results of individual examinations of the effects of gonad type and mass were consistent with our hypotheses. However, upon simultaneous examination of the effects of gonad type and mass, these effects became unclear because of their strong multicollinearity. The compiled data for delimited and non-delimited gonads were biased toward the smaller and larger species, respectively, precluding us from being able to statistically distinguish between these effects. Small species with non-delimited gonads and large species with delimited gonads exist; however, their *T*_*S*_ has not been measured with high temporal resolution thus far. Therefore, additional experiments on these species are required to statistically distinguish between, as well as to better understand, the effects of gonad type and mass on *T*_*S*_.

## RESEARCH QUESTION

Sex change is a well-known phenomenon in plants and animals (Policansky 1982). Female-to-male and male-to-female sex changes have been observed in teleost fishes (Kuwamura et al. 2020). Sex change may span over several days to a few months, depending on the species and direction of sex change. However, the underlying factors impacting the time required for sex change (*T*_*S*_) remain unclear. Given that the time required to produce a new gonad largely determines *T*_*S*_, it is anticipated that there are two factors that may influence *T*_*S*_. First, some sex-changing species retain the gonad of opposite sex with the “delimited” type, where ovarian and testis tissues are separated by a thin cellular wall. The delimited gonad type is considered an adaptation to reduce the amount of new gonads to produce and shorten the *T*_*S*_ (Munday et al. 2010); however, no such quantitative assessment has been conducted thus far. Second, many biological times, such as longevity and gestation periods, are determined by the metabolic rate of animals, which is a function of temperature and body mass (Gillooly et al. 2001, Brown et al. 2004). Consistent with this theory, warmer water causes shorter *T*_*S*_ in sex-changing fish (Black et al. 2005); however, the effect of body mass on *T*_*S*_ requires clarification. As such, this study sought to address two research questions: (1) whether the delimited gonad shortens *T*_*S*_; and (2) whether body mass affects *T*_*S*_.

## VALUE

The ultimate rationale for sex change is to increase fitness (Warner 1975). Sex change does not occur instantly (i.e., it takes several days to a few months), and fish lose breeding opportunities during this period. Under specific ecological and physiological constraints, each species is expected to have evolved to minimize *T*_*S*_. Identifying the factors that determine *Ts* is essential to understand the decision-making process and life history strategies of sex-changing fish. Additionally, identifying such factors affecting *T*_*S*_ would allow researchers to estimate the *T*_*S*_ of various sex-changing species. Such information is useful to induce sex change in commercially important species and enhance seed production in aquaculture (Quinitio et al. 1997, Sato et al. 2018). This information is also essential to estimate the impact of sex-biased harvesting on the reproductive output of wild populations (Sato et al. 2018).

## RELEVANT HYPOTHESIS

This study sought to test two key hypotheses. The first hypothesis is that the *T*_*S*_ of species with delimited gonads is shorter than that of non-delimited gonads. The second hypothesis is that body mass affects *T*_*S*_ through its effect on metabolic rate. More specifically, the metabolic rate *Y* may be expressed by body mass, *M*, as *Y* ∝ *M*^*β*^, where *β* is a metabolic scaling exponent. Biological times, *t*_*B*_, such as gestation period, life-span, and population doubling time, were expressed as *M*/*Y*, that is, *t*_*B*_ ∝ *M*^1−*β*^. As *β* is approximately 0.8–0.9 in fishes (White et al. 2007), we hypothesized that *T*_*S*_ scales with *M*^0.1–0.2^.

From these two hypotheses, we predicted four distinct scenarios. First, if gonad type and <Hbody mass determine *T*_*S*_, the *T*_*S*_ of delimited species is shorter than that of non-delimited species, and the slope in each gonad type is between 0.1 and 0.2 (Figure 1a). Second, if the gonad type alone determines *T*_*S*_, the *T*_*S*_ of delimited species is shorter than that of non-delimited species, and the slope in each gonad type is almost zero (Figure 1b). Third, if body mass alone determines *T*_*S*_, the regression slope is between 0.1 and 0.2, and there is no difference in *Ts* between delimited and non-delimited species (Figure 1c). Fourth, if neither factor affects *T*_*S*_, the slope in each gonad type is almost zero, and there is no difference in *Ts* between delimited and non-delimited species (Figure 1d).

**Figure1.**
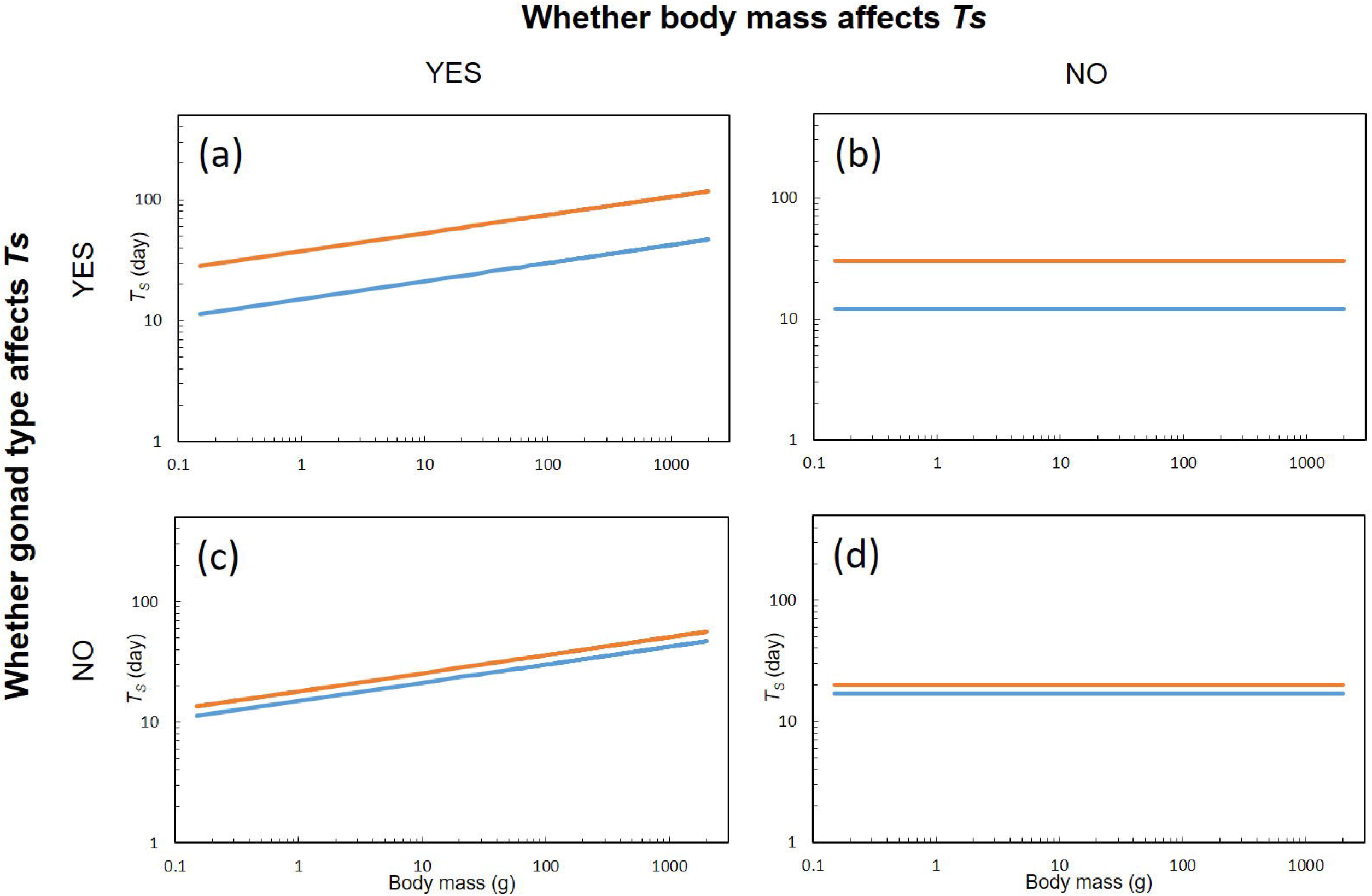
Four possible scenarios of the effects of gonad type and body mass on the time required for sex change (*T*_*S*_). Note that only the scenarios for one sex changing direction are shown, because the scenarios are basically same for both directions. The blue and orange lines represent the expected regression lines for delimited and non-delimited species, respectively. (a) The scenario where both gonad type and body mass determine *T*_*S*_. (b) The scenario where gonad type alone determines *T*_*S*_. (c) The scenario where body mass alone determines *T*_*S*_. (d) The scenario where neither gonad type nor body mass affects *T*_*S*_.

## NEW RESEARCH IDEA

To test these hypotheses, we compiled data on *T*_*S*_ (in days) for as many fish species as possible from the literature. *Ts* was measured as the time between the start of the experiment (e.g., removal of the dominant male, introduction of multiple females into a tank) and when fertilized eggs were first observed. We accepted data from laboratory experiments and the field experiments in which fish were monitored daily. As we are interested in the time from when the physiological switch is turned on and when sex change is complete, we subtracted the period of male–male competition from total periods for the male-to-female sex change. For the female-to-male sex change, the number of females in the mating group may affect the time before the physiological switch is turned on through the change in sex hormone concentrations (Yamaguchi 2016). As such, we excluded experiments where only two females were introduced as a mating group (i.e., all monogamous species and some haremic species), and selected experiments in which the number of females was maximum, when calculating the *T*_*S*_ for each species. We did not model temperature effects because water temperatures were not reported in some studies, and because, for studies reporting water temperatures, their variation among sex-changing species was small (∼25 °C).

All literature compiled in this study reported fish body size as body length. Therefore, we estimated body mass from total length (TL) using Bayesian length-weight relationships (Froese et al. 2014), with the parameters reported in FishBase (Froese & Pauly 2021). Fish size reported as the standard length (SL) or folk length (FL) was converted to TL based on photos available in FishBase or Nakabo (2018). When the size of sex-changing individuals was not reported in the literature, we used the mean length of experimental fish. If fish size was not reported in the literature, we used the size of mature individuals reported in a different study.

To account for the phylogenetic non-independence of species, we first used the phylogenetic generalized least squares (PGLS) method to examine the effects of gonad type and body mass on *T*_*S*_ (see Appendix S1 for details). However, Pagel’s λ was zero in all models, implying that these traits are independent of the given phylogeny, making the ordinary least squares (OLS) method more appropriate (Freckleton et al. 2002). As such, we used the OLS method for subsequent analyses. First, we separately examined the effects of the gonad type and body mass on *Ts*. Thereafter, we examined the concurrent effects of gonad type and body mass on *Ts* using multiple regression analysis. Note that *T*_*S*_ and body mass were log_10_-transformed in all the analyses.

We compiled data on the *T*_*S*_ of 12 and 14 species for female-to-male and male-to-female sex changes, respectively (Table S1). When the effects of gonad type and body mass were examined separately, the results of both were consistent with our hypotheses. The *T*_*S*_ of the species with delimited gonads was shorter than that of non-delimited gonads in both sex-changing directions (female-to-male: *t* = 2.78, *d*.*f*. = 10, *p* = 0.02; male-to-female: *t* = 2.99, *d*.*f*. = 12, *p* = 0.01). The scaling exponent of *Ts* for female-to-male sex change was 0.12 (*t* = 2.51, *d*.*f*. = 10, *p* = 0.03), which was within the range predicted from the metabolic rate (0.1–0.2) (Figure 2a). Although the exponent of *Ts* for male-to-female sex change was slightly larger than the predicted range (slope = 0.32, *t* = 2.46, *d*.*f*. = 12, *p* = 0.03), the 95% confidence interval (CI; 0.04–0.60) included the predicted values (0.1–0.2) (Figure 2b).

**Figure 2.**
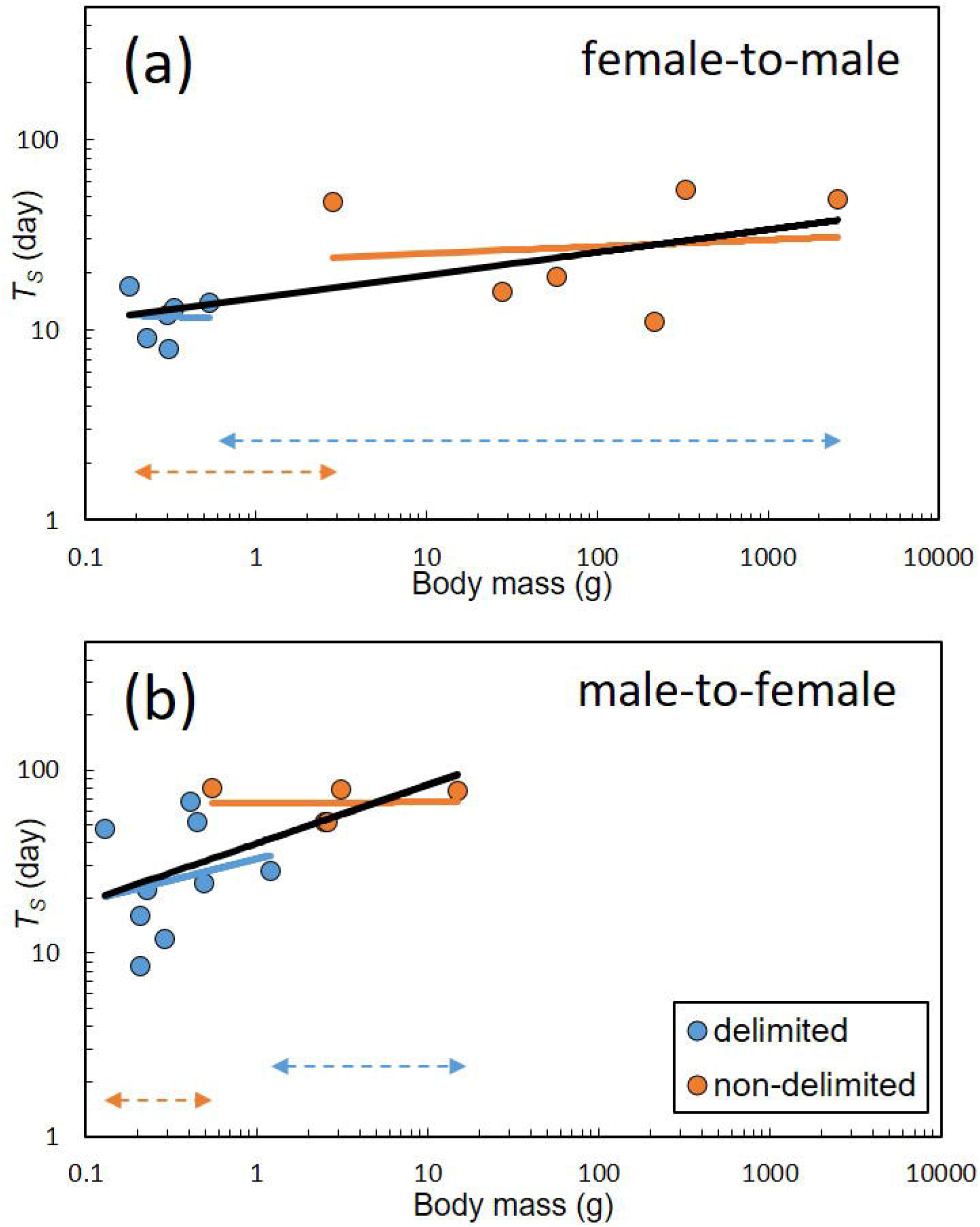
Relationship between body mass and time required for sex change (*T*_*S*_) in (A) female-to-male direction and (B) male-to-female direction. Data are plotted by species. The black solid lines are regression lines for female-to-male (*T*_*s*_ = 14.7 × *mass*^0.12^) and male-to-female sex changes (*T*_*s*_ = 39.8 × *mass*^0.32^). The blue and orange solid lines are regression lines separately calculated for delimited and non-delimited species, respectively. The values of these four slopes are not significantly different from zero (*p* > 0.05). The blue and orange double-headed dotted arrows represent body mass ranges lacking in our dataset for delimited and non-delimited species, respectively.

When the effects of gonad type and mass were examined concurrently, they became unclear in both sex-changing directions. When gonad type was first inputted into the model, the effect of this parameter was significant and the body mass effect was insignificant; however, when body mass was first inputted into the model, the effect of this parameter was significant and the gonad type effect was insignificant (Table 1). These results may be attributable to the strong multicollinearity between gonad type and body mass; the compiled data for delimited and non-delimited gonads were biased toward the smaller (female-to-male, 0.18–0.53 g; male-to-female, 0.13–1.2 g) and larger (female-to-male, 2.8–2561 g; male-to-female, 0.55–15 g) species, respectively (Figure 2; Table S1). This bias precludes the ability to statistically distinguish the effects of gonad type and body mass.

**Table 1.** Effects of body mass, gonad type, and their input orders on the time required for sex change (*T*_*S*_) in each sex-changing direction, as determined by the ordinary least squares multiple regression method.

| First inputted variable | Direction | Source | *df* | *MS* | *F* | *P* |
| --- | --- | --- | --- | --- | --- | --- |
| Body mass | female to male | **Body mass** | **1** | **0.36** | **6.25** | **0.03** |
|  |  | Gonad Type | 1 | 0.05 | 0.90 | 0.37 |
|  |  | Residuals | 9 | 0.06 |  |  |
|  | male to female | **Body mass** | **1** | **0.45** | **6.55** | **0.03** |
|  |  | Gonad Type | 1 | 0.14 | 2.01 | 0.18 |
|  |  | Residuals | 11 | 0.07 |  |  |
| Gonad type | female to male | **Gonad Type** | **1** | **0.40** | **7.04** | **0.03** |
|  |  | Body mass | 1 | 0.01 | 0.11 | 0.75 |
|  |  | Residuals | 9 | 0.06 |  |  |
|  | male to female | **Gonad type** | **1** | **0.57** | **8.36** | **0.01** |
|  |  | Body mass | 1 | 0.01 | 0.20 | 0.66 |
|  |  | Residuals | 11 | 0.07 |  |  |

Significant differences ( $p < 0.05$ ) are in bold; *df*, degrees of freedom; *MS*, mean square. $T_S$ and body mass were $\log_{10}$ -transformed.

Therefore, additional sex-change experiments are required to fill these data gaps (i.e., small species with non-delimited gonads and/or large species with delimited gonads) and to help better understand how *Ts* is determined in fishes.

## HOW TO TACKLE THE QUESTION THROUGH THE PROPOSED NEW IDEA

While some gobiid species (e.g., genera *Coryphopterus, Fusigobius, Gobiodon* and *Paragobiodon*) have non-delimited gonads and small body sizes (Munday et al. 2010, Kuwamura et al. 2020), some sparid species (e.g., genera *Acanthopagrus, Pagrus*, and *Sparus*) have delimited gonads and large body sizes (Buxton & Garratt 1990, Cody & Bortone 1992); their sex change may be observed in large tanks (Kuwamura et al. 2020). Therefore, we propose additional sex-change experiments for these species.

## MOTIVATION

There is a need for additional sex-change experiments, specifically on small species with non-delimited gonads and large species with delimited gonads. Typically, sex-change experiments require specialized techniques that are specific to each species. Thus, we believe that sharing our ideas and data through this “Idea Paper” will guide skilled researchers to focus on the *Ts* of these species.

