## Supplementary material for "Idea Paper: Effects of gonad type and body mass on the time required for sex change in fishes": S1

**SUPPORTING INFORMATION**

**Appendix S1. Phylogenetic generalized least squares (PGLS) analysis**

The PGLS method was used to examine the effects of gonad type and body mass on *T_S_*. This method accounts for the phylogenetic non-independence of species in the dataset (Lavin et al. 2008). We created a phylogenetic tree by combining multiple sources (Maxfield et al. 2012, Gaither et al. 2014, Betancur-R et al. 2017, Sunobe et al. 2017) with an arbitrary branch length (Grafen 1989) using the Mesquite software (Maddison & Maddison 2019) with the PDAP package. We then conducted PGLS analysis using the R software with the “caper” package. For each analysis, we calculated Pagel’s λ (0 ≦ λ ≦ 1) that maximized the model likelihood. Here, λ = 1 means that the trait has evolved according to the given phylogeny and λ = 0 means that the trait is independent of the given phylogeny, and the ordinary least squares (OLS) method is more appropriate (Freckleton et al. 2002). First, we separately examined the effects of gonad type and body mass on *Ts*. Thereafter, we examined the concurrent effects of gonad type and body mass on *Ts*, using multiple regression analysis. As λ was zero in all models, we used the OLS method for subsequent analyses (see main text for the results). Note that *T_S_* and body mass were log_10_-transformed in all the analyses.

**Table S1. Information on time required for sex change (*T_S_*), body mass, and gonad type.**

|  | **Species** | ***T_S_* (day)** | **Body mass (g)** | **Gonad type** | **Reference** |
| --- | --- | --- | --- | --- | --- |
| female-to-male | *Lythrypnus dalli* | 8 | 0.31 | delimited | Reavis & Grober (1999), Black et al. (2005) |
|  | *Trimma kudoi* | 17 | 0.18 | delimited | Manabe et al. (2008) |
|  | *Trimma okinawae* | 12 | 0.30 | delimited | Sunobe & Nakazono (1993) |
|  | *Trimma grammistes* | 9 | 0.23 | delimited | Shiobara (2000) |
|  | *Trimma yanagitai* | 14 | 0.53 | delimited | Sakurai et al. (2008) |
|  | *Trimma caudomaculatum* | 13 | 0.33 | delimited | Tomatsu et al. (2018) |
|  | *Parapercis snyderi* | 47 | 2.8 | non-delimited | Kobayashi et al. (1993) |
|  | *Choerodon schoenleinii* | 49 | 2561 | non-delimited | Sato et al. (2018) |
|  | *Centropyge vroliki* | 16 | 28 | non-delimited | Sakai et al. (2003) |
|  | *Genicanthus lamarck* | 11 | 215 | non-delimited | Suzuki (1979) |
|  | *Genicanthus melanospilos* | 19 | 58 | non-delimited | Hioki et al. (1982) |
|  | *Epinephelus adscensionis* | 55 | 329 | non-delimited | Kline et al. (2011) |
| male-to-female | *Priolepis latifascima* | 48 | 0.13 | delimited | Manabe et al. (2013) |
|  | *Priolepis akihitoi* | 28 | 1.2 | delimited | Manabe et al. (2013) |
|  | *Lythrypnus dalli* | 52 | 0.45 | delimited | Black et al. (2005) |
|  | *Lythrypnus pulchellus* | 12 | 0.29 | delimited | Muñoz-Arroyo et al. (2019) |
|  | *Trimma grammistes* | 22 | 0.23 | delimited | Shiobara (2000) |
|  | *Trimma yanagitai* | 24 | 0.49 | delimited | Sakurai et al. (2008) |
|  | *Trimma caudomaculatum* | 67 | 0.41 | delimited | Tomatsu et al. (2018) |
|  | *Trimma kudoi* | 16 | 0.21 | delimited | Manabe et al. (2008) |
|  | *Trimma okinawae* | 8.5 | 0.21 | delimited | Sunobe & Nakazono (1993) |
|  | *Pseudochromis flavivertex* | 52 | 2.5 | non-delimited | Wittenrich & Munday (2005) |
|  | *Pseudochromis cyanotaenia* | 80 | 0.55 | non-delimited | Wittenrich & Munday (2005) |
|  | *Pseudochromis aldabraensis* | 78 | 3.1 | non-delimited | Wittenrich & Munday (2005) |
|  | *Labroides dimidiatus* | 52 | 2.6 | non-delimited | Kuwamura et al. (2002) |
|  | *Centropyge flavissimus* | 77 | 15 | non-delimited | Hioki & Suzuki (1996) |
